# Antithrombotic Efficacy and Bleeding Risks of Vaccine-Induced Immune Thrombotic Thrombocytopenia Treatments

**DOI:** 10.1101/2024.04.21.589853

**Authors:** H.H.L. Leung, Z. Ahmadi, J. Casey, S. Ratnasingam, S.E. McKenzie, J.S. Perdomo, B.H. Chong

**Author notes:** Correspondence: Beng Chong, Department of Haematology, St. George Clinical Campus, School of Clinical Medicine, Faculty of Medicine and Health, Level 3, 4-10 South Street, Research & Education Centre, Kogarah, New South Wales 2217, Australia.

## Abstract

Current guidelines for treating vaccine-induced immune thrombotic thrombocytopenia (VITT) recommend non-heparin anticoagulants and intravenous immunoglobulin (IVIg). However, the efficacy of these treatments remains uncertain due to a lack of comparative clinical trials or animal studies. A recent study proposed danaparoid and heparin as potential VITT therapies due to their ability to disrupt VITT IgG-PF4 binding. Here, we examined the effects of various anticoagulants (including unfractionated (UF) heparin, danaparoid, bivalirudin, fondaparinux, and argatroban), IVIg, and the FcγRIIa receptor-blocking antibody, IV.3, in relation to VITT pathophysiology. Our investigation focused on VITT IgG-PF4 binding, platelet activation, thrombocytopenia, and thrombosis. Danaparoid, at therapeutic doses, was the sole anticoagulant that reduced VITT IgG-PF4 binding, verified by purified anti-PF4 specific VITT IgG. Low-dose UF heparin (< 2U/mL) augmented VITT IgG binding to PF4 on platelets. While danaparoid and high-dose UF heparin (10 U/mL) inhibited platelet activation, none of the anticoagulants significantly affected thrombocytopenia in our VITT animal model, and all prolonged bleeding time. IVIg and all anticoagulants, except UF heparin, protected VITT mice from thrombosis. Direct FcγRIIa receptor inhibition with IV.3 antibody proved the most effective approach for managing both thrombosis and thrombocytopenia in VITT. Our results underscore the necessity of animal model investigations to inform patient treatment strategies. This study provides compelling evidence for developing FcγRIIa receptor blockers to treat VITT and other FcγRIIa-related thrombotic inflammatory disorders.

**Key points:**

- Non-heparin anticoagulants and IVIg reduce thrombosis *in vivo* by varying degrees whereas heparin exacerbates thrombosis.
- Direct blocking of FcγRIIa receptor is the most effective strategy to treating both thrombosis and thrombocytopenia in VITT.

## Introduction

Adenovirus vector vaccines against SARS-CoV-2 (ChAdOx1 nCoV-19 (Vaxzevria, AstraZeneca) and Ad26.COV2.S (Janssen/Johnson & Johnson) have garnered clinical and public interest due to a rare yet serious complication following vaccination, termed vaccine-induced immune thrombotic thrombocytopenia (VITT). Although the use of the SARS-CoV-2 adenovirus-based vaccines has substantially decreased in high-income countries, it is still widely used in low-medium income countries and has relevance in informing adenovirus-based vaccine development. Additionally, VITT-like immune thromboses following viral infections are increasingly recognised^1-3^ since the advent of VITT.

The pathogenesis of VITT involves the generation of autoantibodies against platelet factor 4 (PF4), with the onset of symptoms appearing 5-30 days post-vaccination. The antigen/antibody immune complex induces platelet and neutrophil activation via FcγIIa^4,5^, resulting in thrombosis. We have previously shown using a humanised VITT animal model that the anti-FcγIIa receptor monoclonal antibody, IV.3, the NETosis inhibitor GSK484 (a reversible peptidylarginine deiminase 4 (PAD4) inhibitor), and genetic ablation of PAD4 in VITT mice led to marked reduction in thrombosis^4^. However, there are currently no inhibitors available in clinical settings that specifically target NETosis or the FcγIIa receptor.

Due to the clinical and pathophysiological similarities between VITT and autoimmune heparin induced thrombocytopenia (HIT), treatment for VITT closely resembles that of HIT. It primarily consists of non-heparin anticoagulants (e.g. direct oral anticoagulants, fondaparinux, danaparoid, and argatroban), intravenous immunoglobulin (IVIg), or less frequently, immune modulatory agents such as steroids or rituximab^6^. At high concentrations, IVIg inhibits the FcγRIIa receptor by non-specific blockage^7^. However, in some cases, VITT patients still develop new major thromboembolic events despite IVIg treatment^8^. Unfractionated (UF) heparin and low-molecular-weight heparin may potentially contribute to disease progression^9^. Nonetheless, a recent study showed that negatively charged anticoagulants like UF heparin and danaparoid disrupt VITT IgG binding to PF4 and inhibit *in vitro* thrombus formation, suggesting their potential effectiveness as VITT treatments^10^. However, *in vitro* findings require validation by *in vivo* studies. Furthermore, the World Health Organization (WHO) Guideline Development Group acknowledged the very low certainty of evidence on treatments for VITT^11,12^. WHO guidelines initially advised against the use of heparin in VITT, then later recommended its use when non-heparin anticoagulants are unavailable^11^. There is a critical need to evaluate and compare the *in vivo* antithrombotic effects of anticoagulants in a relevant mouse model of VITT. These results will assist physicians in selecting the most suitable anticoagulant therapy for VITT and VITT-like conditions, especially in the absence of analogous comparisons from head-to-head clinical trials. While such trials are desirable, their feasibility is challenging. From the wide range of anticoagulants currently used for treatment, our findings identify appropriate anticoagulants for future head-to-head clinical trials, thus improving the feasibility of conducting randomized controlled trials in the future.

In this study, we evaluated the impact of anticoagulants on blocking VITT IgG binding to PF4, platelet activation, thrombocytopenia, and thrombosis. We also compared the efficacy of these anticoagulants with IVIg, using IV.3 as a control. The findings showed that non-heparin anticoagulants inhibited VITT antibody-induced thrombosis to varying degrees, with bivalirudin showing the greatest inhibition. IVIg exhibited comparable efficacy to bivalirudin. However, all anticoagulants prolonged bleeding. Unexpectedly, UF heparin increased *in vivo* thrombosis, contrary to expectations based on *in vitro* results.

## Methods

### Blood samples

Blood was collected with informed consent from patients clinically diagnosed with VITT^13^ and from healthy donors. Six VITT patients were recruited, aged between 46 and 85 (mean age 65), comprising 3 males and 3 females (Table 1). VITT patient samples were positive for laboratory tests (ELISA and ^14^C-serotonin release assay). This study was approved by the Human Research Ethics Committee of South Eastern Sydney Local Health District (17/211 LNR/17/POWH/501).

**Table 1:** VITT patient clinical characteristics. Fib, fibrinogen in g/L; Plt, platelet count x 10^9^/L; D-dimer in mg/L; PE, pulmonary embolism; DVT, deep vein thrombosis.

| VITT | Sex | Site of thrombosis | Platelet count<br>($\times 10^9$/L) | D-Dimer | Fibrinogen |
| --- | --- | --- | --- | --- | --- |
| 1 | M | Portal vein, splenic vein, superior mesenteric vein | 70 | 114 | 3.0 |
| 2 | F | Left central venous sinus | 18 | >20 | 0.6 |
| 3 | M | Bilateral PE, DVT | 21 | >20 | 2.1 |
| 4 | M | Portal vein, superior mesenteric vein | 93 | 56 | 1.9 |
| 5 | F | PE, DVT | 8 | >10 | 1.7 |
| 6 | F | Central venous sinus | 138 | >10 | 2.9 |

### Antibody purification

Total immunoglobulins G (IgG) from healthy donors and patients’ plasma were purified by Protein G Agarose affinity chromatography. Anti-PF4 specific IgG was affinity purified using biotin conjugated PF4 coupled to streptavidin magnetic beads, as previously described^14^. IV.3 monoclonal antibody was produced from hybridoma cells (ATCC clone HB-217) and purified using Protein G Sepharose affinity chromatography. Effector deficient (aglycosylated) IV.3 was used for *in vivo* studies and was prepared as previously described^15^.

### PF4 ELISA

Enzyme-linked immunosorbent assay (ELISA) polystyrene plates (Thermo Scientific) were coated with PF4 (15 μg/mL) with or without heparin (0.1-16 U/mL), danaparoid (0.2-16 U/mL), bivalirudin (2-16 μg/mL), fondaparinux (0.8-16 μg/mL), or argatroban (0.4-16 μg/mL). Concentrations up to 1 U/mL heparin, 0.8U/mL danaparoid, or 1 μg/mL argatroban and fondaparinux were used for affinity purified PF4 specific VITT IgG experiments. ELISA buffer consisted of PBS with 0.05% Tween 20. Fluorescently labelled Alexa Fluor 488 (AF488) (AnaSpec, California)-PF4 was used to quantify the amount of PF4 remaining on the plate following the addition or pre-coating of plates with PF4 and various doses of anticoagulants. VITT patient or control sera (diluted 1:8000) or anti-PF4 specific VITT IgG (5 μg/mL) was added to the ELISA wells. Optical density at 450 nm and fluorescence were measured using a plate reader (Tecan Infinite Pro, Switzerland).

### Serotonin release assay

^14^C serotonin-release assay (^14^C-SRA) was performed as previously described^16^. Briefly, washed donor platelets were incubated with radiolabelled ^14^C, heat inactivated patient’s sera, PF4 (10 μg/mL), and various doses of anticoagulants for 60 min at room temperature while stirring. Reaction was stopped using PBS-EDTA buffer and centrifuged. Radioactivity (counts per minute) of the supernatant was measured using a beta-counter.

### Flow cytometry

Washed platelets were incubated with AF488-labelled PF4 at the optimal concentration (50 μg/mL)^17^ in the absence or presence of heparin (0.1-10 U/mL). AF647-labelled VITT IgG (30 μg/mL) was added to the reaction mixture, incubated at 37 °C for 30 min and analysed by flow cytometry (Fortessa X-20, BD Biosciences, San Jose, CA).

### Mouse model

Double transgenic mice expressing the R^131^ isoform of human FcγRIIa and human PF4 in C57BL/6 background^18^ were injected intravenously with VITT IgG (250 μg/g). Mouse platelets were labelled *in vivo* using anti-CD42c Dylight-649 antibody (1 μg/g, Emfret, Germany). Therapeutic concentrations of anticoagulants followed the 2021 WHO interim guidelines^19^ and South Eastern Sydney Local Health District Medicine Guidelines (heparin (18 U/kg/h), danaparoid (400 U/h), argatroban (120 μg/kg/h), and bivalirudin (0.75 mg/kg))). The anticoagulants were administered via an osmotic pump (Alzet®, CA) implanted the day prior to bleeding assay or administration of VITT IgG. For heparin and danaparoid, a bolus of 0.1 U/g and 0.025 U/g, respectively, was administered 30 min prior to VITT IgG injection or bleeding assay. IVIg (1 g/kg) and IV.3 (1 mg/kg) were administered together with VITT IgG. Mice were bled before IgG administration and at 1 h and 4 h post IgG injection. For thrombus analysis, mice were culled after 4 h, lungs were perfused with PBS followed by formalin, extracted, and imaged using the IVIS Spectrum scanner (Perkin Elmer), as previously described^14,20,21^. Platelet counts were analysed by flow cytometry. Bleeding assays in C57/Bl6 mice were conducted as previously described^22^. Briefly, mice were anaesthetised with a mixture of ketamine (100 mg/kg) and xylazine (10 mg/kg). Mouse tails were amputated 10 mm from the tip then immersed in a 50 mL tube containing warmed (37 °C) saline. Bleeding time was monitored for 20 min. All animal experiments were approved by the UNSW Animal Care and Ethics Committee.

### Statistics

Statistical analysis was performed using GraphPad Prism, Version 9 (GraphPad, La Jolla, CA). Data with normal distribution were analysed by one-way analysis of variance (ANOVA) with Dunnet’s test for multiple comparisons. P values < 0.05 were considered statistically significant.

## Results

### Stability of immobilised PF4 in the presence of anticoagulants

To determine whether anticoagulants interfere with VITT IgG binding to PF4, anticoagulants were added to PF4 coated plates where the anticoagulant concentrations tested encompassed the therapeutic plasma levels (indicated by the dotted brackets in Fig. 1 and Suppl. Fig. 1). We first tested the stability of microtitre plate-immobilized PF4 in the presence of anticoagulants and found that addition of heparin and danaparoid stripped PF4 from the ELISA plates (Suppl. Fig. 1a), whilst argatroban, bivalirudin and fondaparinux had no such effect on plate-bound/immobilized PF4. To minimise PF4 stripping, plates were co-coated with PF4 and various doses of anticoagulants. This approach resolved the stripping effect of danaparoid at all concentrations and of UF heparin at low doses (up to 0.5 U/mL) (Suppl. Fig. 1b). Caution is required when interpreting ELISA data using heparin at concentrations >0.5 U/mL, as well as the effects of danaparoid on binding to PF4 when these anticoagulants are added to the microtitre plates after PF4 coating. This effect is most likely due to the high negative charge of UF heparin and danaparoid.

**Figure 1.**
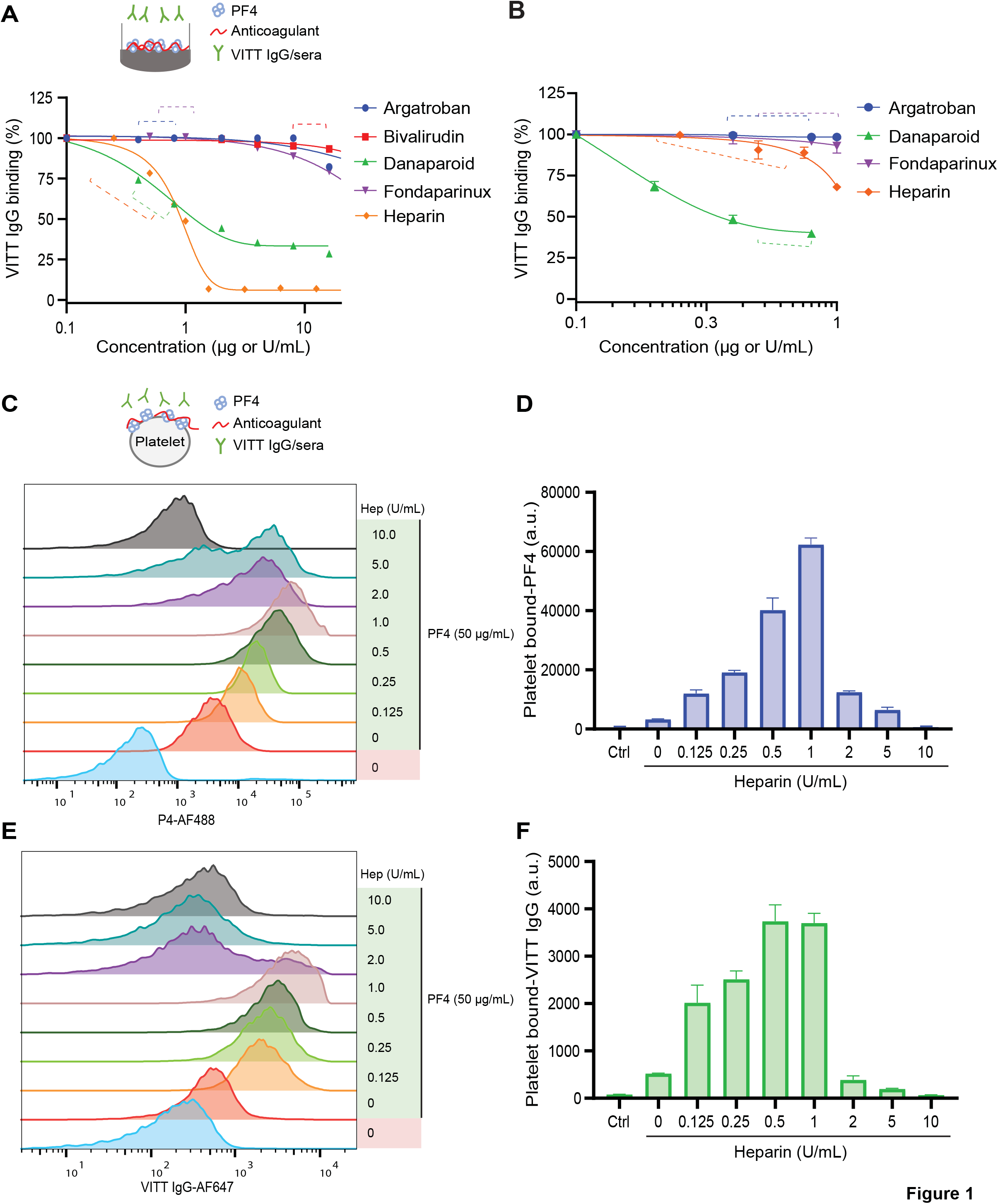
Effect of anticoagulants on VITT IgG binding to microtiter plate-bound PF4 and platelet-bound PF4. Binding of (A) total VITT IgG and (B) anti-PF4 specific VITT IgG to PF4 in the absence or presence of argatroban, bivalirudin, danaparoid, fondaparinux or UF heparin. Data are representative of n=3 VITT patient samples. Dashed lines indicate therapeutic plasma range. Effect of UF heparin on (C-D) Representative histograms and fluorescence quantification (geometric mean, N=3) of PF4 bound to platelets and (E-F) VITT IgG bound to PF4 on platelets, as analysed by flow cytometry. PF4 and VITT IgG were pre-labelled with AF488 and AF647, respectively. Treatment of platelets with VITT IgG in the absence of PF4 and UF heparin was used as a negative control (Ctrl). Hep, heparin. Data shown as mean ± S.D.

### Danaparoid and UF heparin “inhibit” binding of VITT IgG to PF4

Applying the relevant coating steps for the respective anticoagulant, we tested their effect on VITT IgG binding to PF4. At therapeutic doses, danaparoid and UF heparin interfered with VITT IgG binding to PF4 (Fig. 1a). This result corroborates the findings by Singh et al^10^. However, the effect of UF heparin on total VITT IgG binding to PF4 was evident only at the higher end of the therapeutic range (Fig. 1a). To confirm that this inhibitory activity was directed at the specific anti-PF4 antibodies of VITT IgG (Suppl. Fig. 2), we assessed binding using affinity purified anti-PF4 VITT IgG. The data support the finding that danaparoid had a prominent inhibitory effect on specific VITT IgG binding to PF4 at therapeutic concentrations. Notably, UF heparin did not inhibit the binding of anti-PF4 specific VITT IgG to PF4 at low doses (Fig. 1b). The reduction of VITT IgG/PF4 binding observed at a UF heparin concentration of 1 U/mL and higher (Fig. 1a) could be due to the stripping effect of PF4 at these concentrations (Suppl. Fig. 1b), even when UF heparin was added together with PF4 during coating.

### Presence of low dose UF heparin enhances VITT IgG and PF4 binding to platelets

To ascertain the effect of UF heparin on the VITT IgG/PF4 interaction in a more physiological system, we assessed the effect of UF heparin on VITT IgG binding to platelet-bound PF4. UF heparin is known to increase PF4 binding to platelets at low doses^23^, therefore we first incubated washed platelets with labelled PF4 in the absence and presence of increasing amounts of UF heparin. We found that low doses of UF heparin enhanced PF4 association with platelets (Fig. 1c-d). This supports the findings of a previous study that showed UF heparin increased PF4 binding in a dose-dependent manner^14^. Interestingly, VITT IgG bound to PF4 on platelets in an analogous manner and reached a peak at UF heparin concentrations between 0.5-1 U/mL (Fig. 1e-f). UF heparin doses ≥ 2 U/mL substantially decreased binding of VITT IgG (Fig. 1e-f). Although the presence of UF heparin enhances PF4 binding to platelets, with a maximal 18-fold increase (Fig. 1c-d), VITT IgG binding to platelets increases to a lesser extent (7-fold increase, Fig. 1e-f).

### Danaparoid inhibits VITT IgG-induced platelet activation

Next, we investigated the effect of anticoagulants on platelet activation using serotonin release assay, with a cut-off set at 20% (for negative results)^16^. Our findings indicate that only danaparoid completely blocked platelet activation, while increasing concentrations of UF heparin resulted in reduced activation (Fig. 2). Platelet activation was fully inhibited by UF heparin only at a concentration of 10 U/mL, which exceeds the therapeutic range.

**Figure 2.**
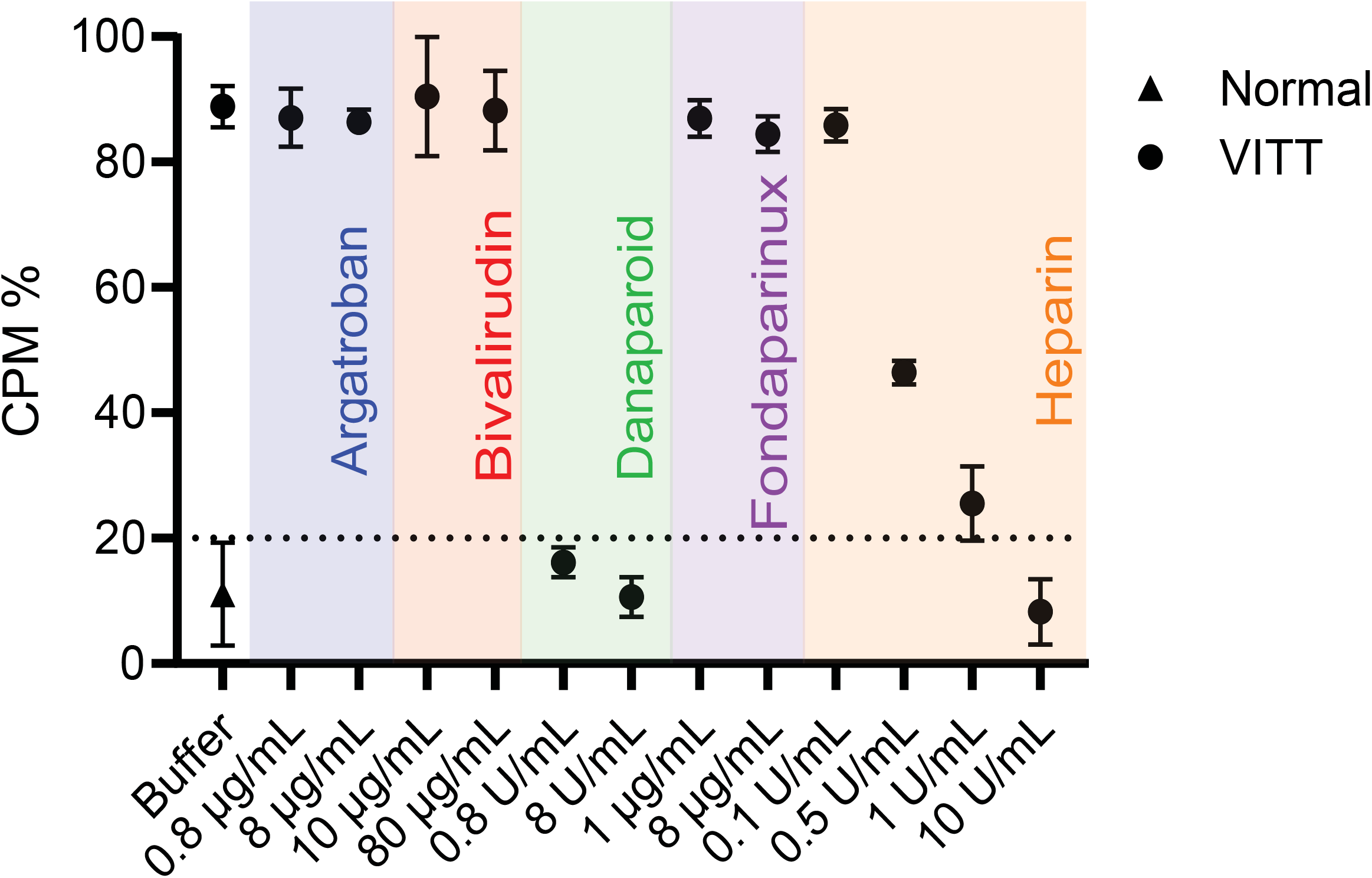
Danaparoid and UF heparin block platelet activation by VITT IgG. Serotonin release assay shows that VITT-induced platelet activation is inhibited in the presence of therapeutic (0.8 μg/mL) and high dose (8 μg/mL) danaparoid and high dose UF heparin (10 U/mL) but not argatroban (0.8 and 8 μg/mL), bivalirudin (10 and 80 μg/mL), fondaparinux (1 and 8 μg/mL) or low dose UF heparin (0.1, 0.5 and 1 U/mL). Data shown as mean ± S.D. Data representative of n=3 VITT patient samples. CPM, counts per minute.

### VITT IgG-induced thrombocytopenia is not improved by anticoagulants

To assess how these findings translate to the clinical manifestations of thrombosis and thrombocytopenia in VITT, anticoagulants were administered in VITT mice at human equivalent doses. The protocol is summarised in Fig. 3a. Mice treated with bivalirudin, argatroban, danaparoid, or UF heparin were not protected from thrombocytopenia (Fig. 3b), while IVIg (Fig. 3c) and IV.3 (Fig. 3d) treatments provided minimal and substantial platelet protection, respectively. Overall, these results are consistent with the mechanisms of anticoagulants, i.e. inhibition of the coagulation cascade rather than blockage FcγRIIa-mediated cell activation. Nevertheless, these findings underscore the value of *in vivo* observations, as anticoagulants that affect VITT antibody binding and antibody-induced platelet activation *in vitro* (danaparoid and UF heparin, Figs. 1 and 2) did not prevent thrombocytopenia in our mouse model. Conversely, IVIg and IV.3 which interfere with FcγRIIa cell signalling by nonspecific and specific blockade of FcγRIIa, respectively, protected platelets from clearance. These results suggest that thrombocytopenia in VITT is due to clearance of platelets coated with VITT IgG immune complexes, rather than peripheral consumption of activated platelets by extensive thrombosis.

**Figure 3.**
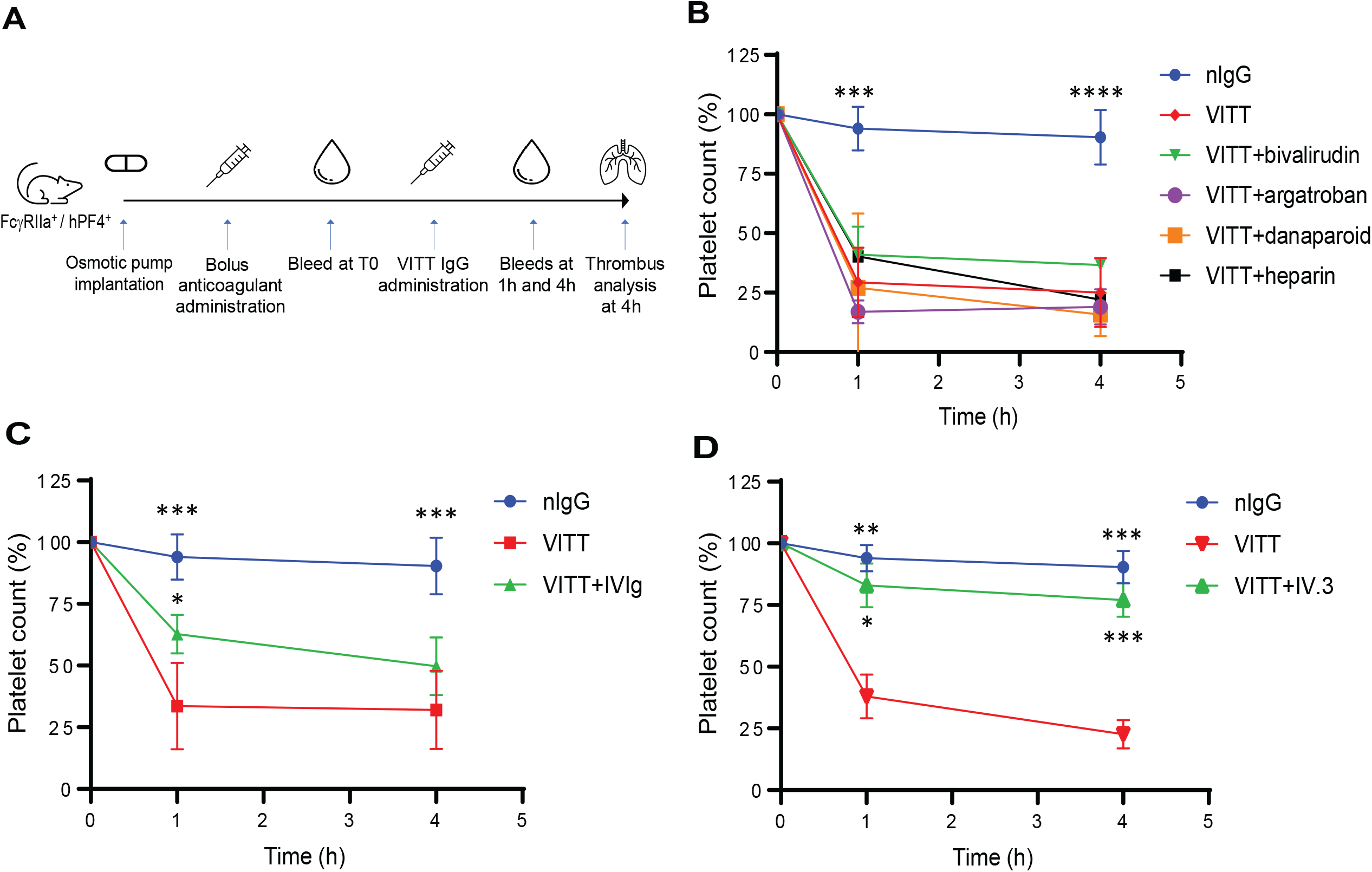
IVIg and IV.3 provide partial and full protection, respectively, from platelet destruction *in vivo*. (A) Study timeline of animal experiment. The VITT condition was recreated in FcγRIIa^+^/hPF4^+^ transgenic mice by administration of VITT IgG. Anticoagulants were administered via osmotic pump and/or intravenous injection, as described in the Methods section. Platelet counts were assessed in mice following treatment with (B) anticoagulants (argatroban, bivalirudin, fondaparinux, danaparoid or UF heparin), (C) IVIg, and (D) aglycosylated IV.3. * *p* < 0.05; ** *p* < 0.01; *** *p* < 0.001; **** *p* < 0.0001, relative to VITT IgG. Data shown as mean ± S.D.

### Anticoagulants, except UF heparin, inhibit VITT IgG-induced thrombosis

Despite the lack of protection against thrombocytopenia, treatment with bivalirudin, argatroban, or danaparoid led to significant but varying degrees of reduction in clot formation, as assessed by thrombus embolization in VITT mouse lungs. Bivalirudin demonstrated the strongest inhibition, comparable to IVIg. In contrast, UF heparin treatment (18U/kg/h) significantly exacerbated thrombosis (Fig. 4a-b). This result aligns with the VITT IgG-PF4-platelet binding data depicted in Fig. 1c-d, indicating that therapeutic doses of UF heparin enhances PF4 and consequently VITT IgG binding to platelets. The blockage of FcγRIIa with aglycosylated IV.3 provided the greatest reduction in thrombosis. Statistically, however, IVIg showed no significant difference compared to anticoagulants, except UF heparin.

**Figure 4.**
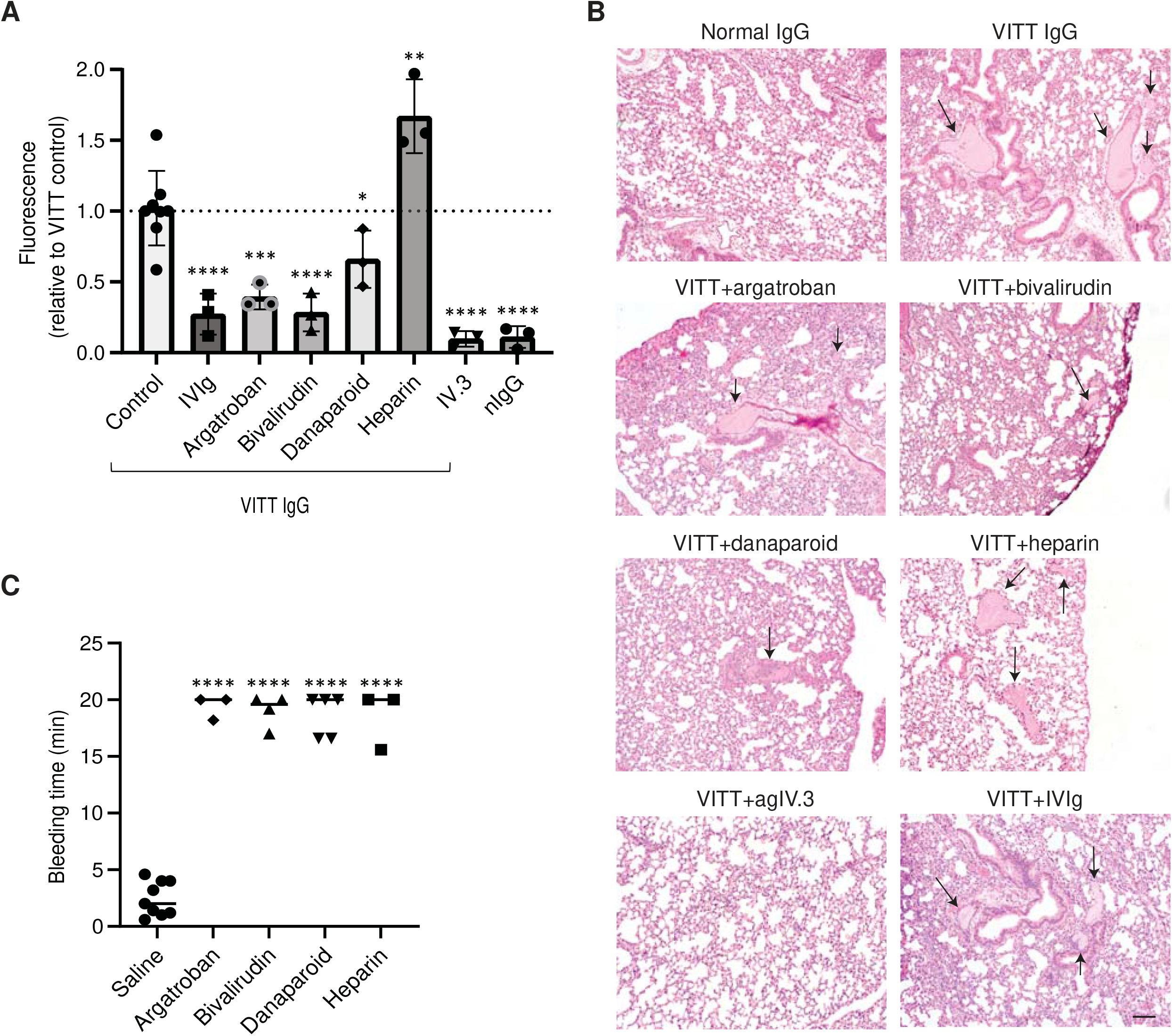
Non-heparin anticoagulants, IVIg and IV.3 inhibit but UF heparin enhances VITT-induced thrombosis in mice. (A) Mouse lungs from FcγRIIa^+^/hPF4^+^ mice treated with VITT IgG, with or without anticoagulants, were extracted and analysed for thrombosis by fluorescence using the IVIS Spectrum CT. (B) Representative images of H&E staining of lung sections. Images were acquired with a 10x objective using an inverted Olympus CKX53 microscope. Arrows indicate clots. Scale bar, 50 μm. (C) Mice treated with UF heparin, bivalirudin, argatroban, or danaparoid at therapeutic concentrations had prolonged bleeding times compared to saline control. Bleeding time was measured over 20 min. * *p* < 0.05; ** *p* < 0.01; *** *p* < 0.001; **** *p* < 0.0001, relative to VITT IgG (A) or saline (C). nIgG, normal IgG. Data shown as mean ± S.D.

A well-known limitation of anticoagulant treatment is the risk of serious bleeding^24^. To examine the impact of anticoagulants on haemostasis, tail bleeding assays were conducted on mice following treatment with clinically relevant doses. Mice treated with UF heparin, bivalirudin, argatroban, or danaparoid all experienced significant prolongation of bleeding times (Fig. 4c). Our *in vivo* data collectively highlights the necessity for the development of specific anti-FcγRIIa receptor inhibitors. Such inhibitors could offer more targeted and effective treatment options to prevent both thrombocytopenia and thrombosis without the bleeding side effects associated with traditional anticoagulants.

### Discussion

Despite the lack of comprehensive data on the efficacy of different treatment regimens for VITT derived from clinical trials or animal studies, treatment guidelines for VITT have been established based on its clinical and pathophysiological similarities to HIT. Experts in the field recognise that many questions remain unanswered, including whether VITT patients should avoid heparin use and which treatments are likely to yield the best clinical outcomes^9^. This study evaluated in detail the effect of anticoagulants *in vitro* and in a VITT mouse model, with the latter also compared with IVIg. Our findings indicate that current treatments such as non-heparin anticoagulants and IVIg effectively block thrombosis in our mouse model, albeit at the cost of prolonged bleeding time in the case of anticoagulants. Importantly, these observations support the notion that heparin should not be used as a treatment for VITT^10^, as evidenced by our data showing that UF heparin can enhance VITT antibody-induced thrombosis. This phenomenon is likely attributable to the presence of antibodies in VITT patients that bind both PF4-heparin and PF4^25,26^. Similarly, low dose UF heparin (0.2 U/mL) was shown to enhance platelet aggregation in 1 out of 5 VITT patients tested^8^. Several VITT studies have suggested accelerated disease progression following treatment with UF heparin or low-molecular-weight heparin^13,27^. Moreover, a large prospective study reported a higher mortality rate (20%) among VITT patients treated with heparin compared to those receiving non-heparin anticoagulants (16%)^28^. Our data further demonstrates that UF heparin enhances PF4 and VITT IgG binding to platelets, thereby amplifying platelet activation and thrombosis in VITT, as observed *in vivo*. Similarly, UF heparin may have the capacity to increase binding of immune complexes to neutrophils, potentially triggering neutrophil activation, release of neutrophil extracellular traps (NETs), and exacerbating the thrombotic process. Consequently, the administration of UF heparin to patients with VITT should be given with great caution. Although the sample size of this study is limited, our findings are likely applicable to VITT in general due to the clonotypic nature of VITT antibodies^29^.

Similar to recent findings^10^, we show that only danaparoid and high dose UF heparin effectively inhibit platelet activation *in vitro*. We also observe that strong negatively charged anticoagulants, such as UF heparin, not only block the binding of VITT IgG to immobilised PF4 but also interfere with the binding stability of positively charged PF4 to the ELISA plate surface. This observation underscores the importance of exercising caution when interpreting results involving high supra-therapeutic doses of UF heparin. The interaction between heparin and PF4 may introduce potential confounding factors in the analysis, highlighting the necessity for careful consideration in experimental design and result interpretation.

We expanded our investigation to explore the efficacy of IVIg and various anticoagulants in blocking thrombosis *in vivo*, thereby enhancing the clinical relevance of our study. We found that IVIg was most effective at improving platelet counts and reducing thrombosis, and unlike anticoagulants, IVIg does not prolong bleeding time. Although IVIg is recommended over other current VITT treatments, potential side effects of IVIg should be noted. IVIg is derived from pooled plasma of 3,000-100,000 healthy donors^30^ and very high concentrations are required for treatment to achieve the desired effect of non-specific inhibition of FcγRIIa. Aggregates of IVIg can be found in batches of IVIg and can cause paradoxical prothrombotic effects, platelet aggregation^31^, and potential thromboembolism^32,33^. In contrast, IV.3 requires doses 1000 times less than IVIg to achieve specific FcγRIIa inhibition to a greater extent. Nonetheless, reports suggest that high-dose IVIg in VITT patients can increase platelet counts and inhibit the generation of procoagulant platelets^34^, thus making it a recommended treatment option for VITT patients, considering the potential clinical benefits outweigh the uncommon but serious complications. Another promising treatment avenue for VITT being explored is a deglycosylated form of an anti-PF4 antibody (1E12) that recognizes overlapping epitopes on PF4 as VITT IgG^35^. This approach has shown promise in inhibiting VITT-induced cell activation and thrombosis *in vitro*.

In summary, our findings provide evidence supporting the necessity for developing more targeted therapies to mitigate unwanted bleeding side effects. UF heparin should not be considered as a first-line VITT treatment due to its potential prothrombotic complications, as shown by the results of this study. The prothrombotic effects of UF heparin likely stem from the enhancement of VITT IgG binding to PF4-heparin complexes on cell surfaces. While current anticoagulant treatments effectively block thrombosis in VITT, there is a need for more targeted therapies to address the associated bleeding side effects. Given the ongoing clinical relevance of VITT, particularly in regions where adenovirus-based vaccines are still widely used, our study serves as a valuable guide for anticoagulant selection in the treatment of VITT, VITT-like thromboses, HIT (particularly autoimmune HIT) and potentially other immune-thromboses^36^. The efficacy of FcyRIIa blockade may lead to improved treatment options for these potentially fatal conditions. Our study provides a comprehensive understanding of how these therapeutic agents impact thrombotic events in a physiological context and sheds light on the potential implications for treatment strategies and patient care in the context of VITT and related conditions.

## Acknowledgements

The authors would like to thank Drs Shiying Zheng, Rose Wong and Feng Yan for assistance with patient samples. *In vivo* imaging data presented in this study was acquired at the Mark Wainwright Analytical Centre (MWAC) of UNSW Sydney, which is in part funded by the Research Infrastructure Program of UNSW. This study was funded by grants from the National Health and Medical Research Council (APP1052616 to B.H.C. and APP2030328 to H.H.L.L. and B.H.C.) and the New South Wales Health Government Office for Health and Medical Research (Senior Researcher Grant to B.H.C. and Cardiovascular Collaborative Grant to H.H.L.L. and B.H.C.).

## Authorship Contributions

H.H.L.L. conceived the study, designed and performed experiments, analyzed and interpreted data, and wrote the manuscript; Z.A. performed experiments and analyzed data; S.McK. provided FcγRIIa^+^/hPF4^+^ transgenic mice and critically reviewed the manuscript; S.R. and J.C. provided clinical input, and critically reviewed the manuscript; J.P. designed and performed experiments, supervised the study, provided intellectual input, and critically reviewed manuscript; and B.H.C. conceived and supervised the study, provided intellectual input, and critically reviewed the manuscript.

## References

1 Greinacher, A. et al. Platelet-activating anti-PF4 antibodies mimic VITT antibodies in an unvaccinated patient with monoclonal gammopathy. Haematologica 107, 1219–1221, doi:10.3324/haematol.2021.280366 (2022).

2 Uzun, G. et al. Cerebral venous sinus thrombosis and thrombocytopenia due to heparin-independent anti-PF4 antibodies after adenovirus infection. Haematologica, doi:10.3324/haematol.2023.284127 (2023).

3 Warkentin, T. E. et al. Adenovirus-Associated Thrombocytopenia, Thrombosis, and VITT-like Antibodies. N Engl J Med 389, 574–577, doi:10.1056/NEJMc2307721 (2023).

4 Leung, H. H. L. et al. NETosis and thrombosis in vaccine-induced immune thrombotic thrombocytopenia. Nat Commun 13, 5206, doi:10.1038/s41467-022-32946-1 (2022).

5 Greinacher, A. et al. Insights in ChAdOx1 nCoV-19 vaccine-induced immune thrombotic thrombocytopenia. Blood 138, 2256–2268, doi:10.1182/blood.2021013231 (2021).

6 Klok, F. A., Pai, M., Huisman, M. & Makris, M. Vaccine-induced immune thrombotic thrombocytopenia. Lancet Haematol 9, e73–80 (2022).

7 Warkentin, T. E. High-dose intravenous immunoglobulin for the treatment and prevention of heparin-induced thrombocytopenia: a review. Expert Rev Hematol 12, 685–698, doi:10.1080/17474086.2019.1636645 (2019).

8 Tiede, A. et al. Prothrombotic immune thrombocytopenia after COVID-19 vaccination. Blood 138, 350–353, doi:10.1182/blood.2021011958 (2021).

9 Arepally, G. M. & Ortel, T. L. Vaccine-induced immune thrombotic thrombocytopenia: what we know and do not know. Blood 138, 293–298 (2021).

10 Singh, A. et al. The interaction between anti-PF4 antibodies and anticoagulants in vaccine-induced thrombotic thrombocytopenia. Blood 139, 3430–3438 (2022).

11 World Health Organization TTS Guideline Development Group. (ed World Health Organization) (2023).

12 Organization, W. H. Vol. WHO/2019-nCoV/TTS/2021.1 (ed World Health Organization) (2021).

13 Schultz, N. H. et al. Thrombosis and Thrombocytopenia after ChAdOx1 nCoV-19 Vaccination. N Engl J Med 384, 2124–2130, doi:10.1056/NEJMoa2104882 (2021).

14 Leung, H. et al. NETosis and thrombosis in vaccine-induced immune thrombotic thrombocytopenia. Nature Commun 13, 5206 (2022).

15 Perdomo, J. et al. Neutrophil activation and NETosis are the major drivers of thrombosis in heparin-induced thrombocytopenia. Nature Commun 10, 1322, doi:10.1038/s41467-019-09160-7 (2019).

16 Sheridan, D., Carter, C. & Kelton, J. G. A diagnostic test for heparin-induced thrombocytopenia. Blood 67, 27–30 (1986).

17 Rauova, L. et al. Role of platelet surface PF4 antigenic complexes in heparin-induced thrombocytopenia pathogenesis: diagnostic and therapeutic implications. Blood 107, 2346–2353, doi:10.1182/blood-2005-08-3122 (2006).

18 Reilly, M. P. et al. Heparin-induced thrombocytopenia/thrombosis in a transgenic mouse model requires human platelet factor 4 and platelet activation through FcgammaRIIA. Blood 98, 2442–2447, doi:10.1182/blood.v98.8.2442 (2001).

19 World Health Organization. Guidance for clinical case management of thrombosis with thrombocytopenia syndrome (TTS) following vaccination to prevent coronavirus disease (COVID-19). (2021).

20 Leung, H. H. L. et al. Inhibition of NADPH oxidase blocks NETosis and reduces thrombosis in heparin-induced thrombocytopenia. Blood Adv 5, 5439–5451, doi:10.1182/bloodadvances.2020003093 (2021).

21 Perdomo, J. et al. Neutrophil activation and NETosis are the major drivers of thrombosis in heparin-induced thrombocytopenia. Nat Commun 10, 1322, doi:10.1038/s41467-019-09160-7 (2019).

22 Liu, Y., Jennings, N. L., Dart, A. M. & Du, X. J. Standardizing a simpler, more sensitive and accurate tail bleeding assay in mice. World J Exp Med 2, 30–36, doi:10.5493/wjem.v2.i2.30 (2012).

23 Jaax, M. E. et al. Complex formation with nucleic acids and aptamers alters the antigenic properties of platelet factor 4. Blood 122, 272–281, doi:10.1182/blood-2013-01-478966 (2013).

24 Cundiff, D. K. A systematic review of Cochrane anticoagulation reviews. Medscape J Med 11, 5 (2009).

25 Althaus, K. et al. Antibody-mediated procoagulant platelets in SARS-CoV-2-vaccination associated immune thrombotic thrombocytopenia. Haematologica 106, 2170–2179, doi:10.3324/haematol.2021.279000 (2021).

26 Greinacher, A. et al. Thrombotic Thrombocytopenia after ChAdOx1 nCov-19 Vaccination. N Engl J Med 384, 2092–2101 (2021).

27 Scully, M. et al. Pathologic Antibodies to Platelet Factor 4 after ChAdOx1 nCoV-19 Vaccination. N Engl J Med 384, 2202–2211, doi:10.1056/NEJMoa2105385 (2021).

28 Pavord, S. et al. Clinical Features of Vaccine-Induced Immune Thrombocytopenia and Thrombosis. N Engl J Med 385, 1680–1689, doi:10.1056/NEJMoa2109908 (2021).

29 Wang, J. J. et al. Vaccine-induced immune thrombotic thrombocytopenia is mediated by a stereotyped clonotypic antibody. Blood 140, 1738–1742, doi:10.1182/blood.2022016474 (2022).

30 Kazatchkine, M. D. & Kaveri, S. V. Immunomodulation of autoimmune and inflammatory diseases with intravenous immune globulin. N Engl J Med 345, 747–755, doi:10.1056/NEJMra993360 (2001).

31 Pollreisz, A. et al. Intravenous immunoglobulins induce CD32-mediated platelet aggregation in vitro. Br J Dermatol 159, 578–584, doi:10.1111/j.1365-2133.2008.08700.x (2008).

32 Yu, C. F. et al. Acute pulmonary embolism caused by highly aggregated intravenous immunoglobulin. Vox Sang 110, 27–35, doi:10.1111/vox.12307 (2016).

33 Ammann, E. M. et al. Intravenous immune globulin and thromboembolic adverse events in patients with hematologic malignancy. Blood 127, 200–207, doi:10.1182/blood-2015-05-647552 (2016).

34 Uzun, G. et al. The use of IV immunoglobulin in the treatment of vaccine-induced immune thrombotic thrombocytopenia. Blood 138, 992–996, doi:10.1182/blood.2021012479 (2021).

35 Vayne, C. et al. The deglycosylated form of 1E12 inhibits platelet activation and prothrombotic effects induced by VITT antibodies. Haematologica 107, 2445–2453, doi:10.3324/haematol.2021.280251 (2022).

36 Perdomo, J. & Leung, H. H. L. Immune Thrombosis: Exploring the Significance of Immune Complexes and NETosis. Biology (Basel) 12, doi:10.3390/biology12101332 (2023).

