## Supplementary Figures for "Antithrombotic Efficacy and Bleeding Risks of Vaccine-Induced Immune Thrombotic Thrombocytopenia Treatments"

**A**

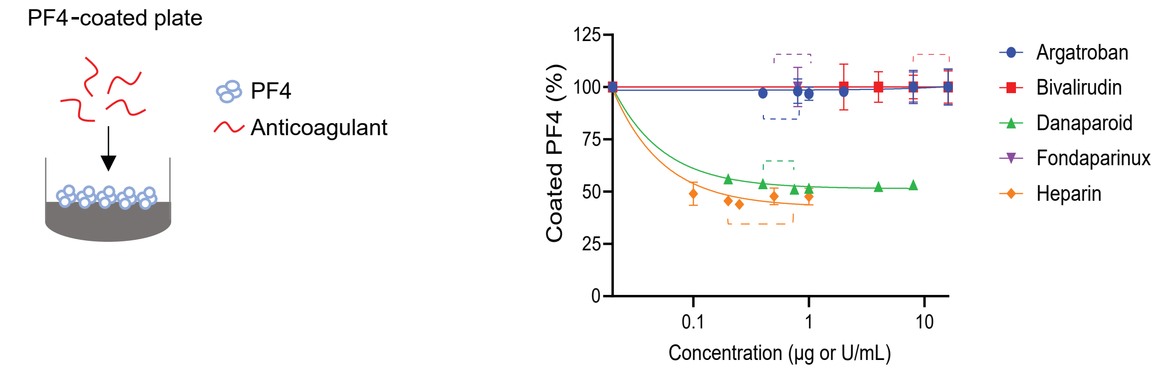


**B**

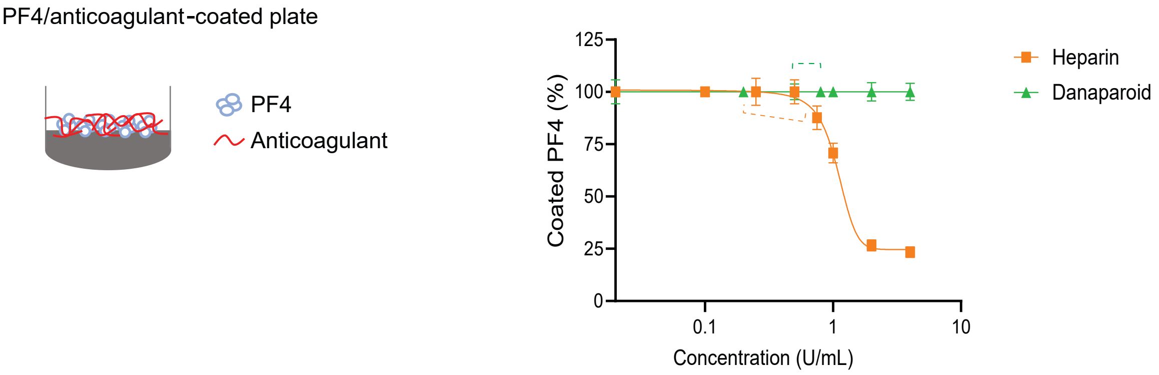


**Negatively charged anticoagulants interfere with PF4 bound to ELISA plates.** (**A**) Addition of argatroban, bivalirudin, or fondaparinux up to 16 ug/mL have no effect on plate-bound PF4, while low doses of UF heparin (from 0.1 U/mL) or danaparoid (from 0.25 U/mL) strip off PF4 from the plate. (**B**) Co-incubation of PF4 and UF heparin or PF4 and danaparoid on ELISA plates stabilises plate-bound PF4. Dashed lines indicate therapeutic plasma level range. Data shown as mean ± S.D.

**Supplementary Figure 1**

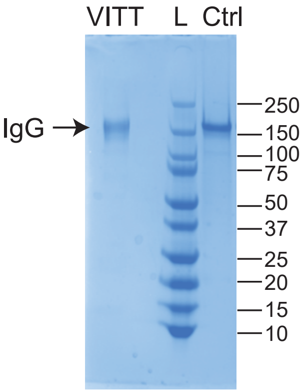


**Affinity purified anti-PF4 IgG from VITT patient sera.** Non-reducing SDS gel showing affinity purified anti PF4 IgG from VITT patient (VITT). Commercial monoclonal IgG (Ctrl) used as a comparison. L indicates ladder.

Supplementary Figure 2
